# Higher-order effects, continuous species interactions, and trait evolution shape microbial spatial dynamics

**DOI:** 10.1101/2021.02.02.429382

**Authors:** Anshuman Swain, Levi Fussell, William F Fagan

**Author notes:** contributed equally (Anshuman Swain, Levi Fussell). Corresponding author: Anshuman Swain, **Email**.

## Abstract

The assembly and maintenance of microbial diversity in natural communities, despite the abundance of toxin-based antagonistic interactions, presents major challenges for biological understanding. A common framework for investigating such antagonistic interactions involve cyclic dominance games with pairwise interactions. The incorporation of higher-order interactions in such models permits increased levels of microbial diversity, especially in communities where antibiotic producing, sensitive, and resistant strains co-exist. However, most such models involve a small number of discrete species, assume a notion of pure cyclic dominance, and focus on low mutation rate regimes, none of which well represents the highly interlinked, quickly evolving, and continuous nature of microbial phenotypic space. Here, we present an alternative vision of spatial dynamics for microbial communities based on antagonistic interactions—one in which a large number of species interact in continuous phenotypic space, are capable of rapid mutation, and engage in both direct and higher-order interactions mediated by production of and resistance to antibiotics. Focusing on toxin production, vulnerability, and inhibition among species, we observe highly divergent patterns of diversity and spatial community dynamics. We find that species interaction constraints (rather than mobility) best predict spatiotemporal disturbance regimes, whereas community formation time, mobility, and mutation size best explain patterns of diversity. We also report an intriguing relationship among community formation time, spatial disturbance regimes, and diversity dynamics. This relationship, which suggests that both higher-order interactions and rapid evolution are critical for the origin and maintenance of microbial diversity, has broad-ranging links to the maintenance of diversity in other systems.

**Significance Statement:** Persistently diverse microbial communities are one of biology’s great puzzles. Using a novel continuous trait space modeling framework that accommodates high mutation rates, elevated species richness, and direct and higher-order antagonistic species interactions, we find that two parameters characterizing mutation size and mobility best explain patterns of microbial diversity. Moreover, community formation time (the duration of the transient phase in community assembly) provides an unexpectedly clear guide to the diversity profiles of the resulting communities. These discoveries showcase how complex, antagonistic interactions mediated by the production of, inhibition of, and vulnerability to toxins (antibiotics) can shape microbial communities, allowing for extraordinarily high levels of diversity and temporal persistence.

## 1. Introduction

Understanding the origin and maintenance of species diversity is a long-standing biological question, and understanding diversity through the lens of species interactions has been deemed key (1–5). Natural communities possess high species diversity, often much higher than expected from a ‘simple’ Darwinian interpretation of species interactions (6). Extraordinary levels of diversity exist in several macroscopic systems (e.g., coral reefs, tropical forests), but remarkable levels of diversity also exist in microscopic systems, among microbial communities. High levels of diversity in microbial systems are especially intriguing given the ubiquity of antagonistic interactions among microbes, such as the production of antibiotics/toxins against one another (7–11).

Theoretical and empirical studies have explored microbial interactions with the goal of understanding the mechanisms of species coexistence in hyper-diverse microbial systems. Major examples include models with: (**A**) mutualistic and cross-feeding interactions (12); (**B**) resource-based competitive interactions (13); (**C**) a combination of mutualistic and competitive interactions (13–16); (**D**) stochastic spatio-temporal processes (17,18); and (**E**) antagonistic interactions due to toxin production (6, 19, 20).

Previous studies focusing on the mutualistic, cross-feeding and resource-based competitive interactions (points **A-C** above) have demonstrated that positive interactions arising from resource exchange can affect key criteria governing the relationship between community diversity and stability. Examples of such criteria are that species richness is inversely related to the strength of random pairwise interactions in stable communities (2) and communities constructed with random pairwise interactions feature an upper bound on diversity (21–23). Additional key findings from the resource-based framework are that cooperative (metabolic) interactions can arise spontaneously in complex microbial communities (24) and that a cooperative versus competitive dichotomy of microbial communities may exist, depending on conditions (25). Sometimes, the interactions arising from resource exchange allow systems to bypass classic stability-diversity relationships (13, 14). Collectively, these results have tremendously improved our understanding of the factors influencing the development and persistence of microbial community diversity. However, resource-based perspectives fail to account for a major class of well-known antagonistic interactions that exist among microbes, mediated through bacterial toxins or antibiotics (also termed as bacteriocins and related compounds). Toxin-based interactions are ubiquitous in both natural microbial communities and controlled settings (7–10), are found in all known phyla of bacteria, come in diverse forms, and play a central, yet usually underappreciated role in shaping microbial community structure (11).

Here, we focus on modeling microbial communities structured by antagonistic interactions among antibiotic/toxin producing, sensitive and resistant species. Prior research in this area has relied on the framework of cyclic dominance (also termed non-transitive systems or rock-paper-scissors systems) and related techniques in evolutionary game theory (7, 19, 20, 26–28). Early models using cyclic dominance explored community stabilization, increased co-existence, and maintenance of diversity in spatially structured environments (19, 26, 29, 30). However, because these models focused on pairwise interactions, they generally did not account for other kinds of interactions that occur when intermixed species coexist at very small spatial scales (6, 9, 10, 31–33). For example, real microbial species are routinely involved in multi-species interaction systems including so-called ‘higher-order interactions’ in which the interactions between two species can be modulated by other species (34, 35). One simple example of such a behavior occurs when an antibiotic produced by one species, that inhibits the growth of a competing (sensitive) species, can be attenuated by a third species that can degrade the antibiotic (20, 36, 37). Here, the third species modulates the interaction between the antibiotic-producing and antibiotic-sensitive species, without impacting either of them directly (34), thus creating a higher-order interaction.

Models using cyclic dominance with higher-order interactions for a small number of static (i.e., non-evolving) species have demonstrated increased stability and diversity in both spatially structured and well-mixed communities (6, 19, 20, 26, 29, 38), and findings from such models have been verified empirically (20). In contrast, horizontal gene transfer and mutation blur the boundaries among microbial strains and species, reflecting the continuous nature of phenotypes in microbial assemblages. Scaling up analytic models of cyclic dominance to species-rich scenarios is not possible because the incorporation of more than a few discrete species makes the dynamics extremely complex and hard to interpret using equation-based approaches (20).

Despite its prominence in modeling studies, pure cyclic dominance, in which there is a ‘closed’ loop of dominance, is rare in nature (39–42). In contrast, rapid evolution routinely leads to model systems featuring semi-cyclic or non-cyclic patterns in which higher-order interactions, such those involving antibiotic producing, sensitive, and resistant species, are common (35, 40). Therefore, the notion of pure cyclic dominance used in earlier studies of antagonistic microbial dynamics needs to be supplemented with evolving, ‘mixed’ patterns of dominance, in order to better model microbial community interactions.

Prior studies of community-level eco-evolutionary dynamics, which have explored the evolution of species interactions abstractly or the specific traits that determine such interactions (43–45), focused on niche-based food web models (i.e., those involving resource consumption) rather than the non-resource based antagonistic interactions that characterize our work. Even when studies did investigate antagonistic interactions and resultant species dynamics, they emphasized very slow mutation regimes and assumed a separation of ecological and evolutionary timescales (46–48). Such assumptions do not work very well for microbial systems, where high mutation rates and extensive horizontal gene transfers are commonplace, creating an overlap of ecological and evolutionary timescales (47–50). These eco-evolutionary feedbacks can affect the nature of trait evolution (48) and even destabilize species interactions (51). Recent models exploring these fast-evolving regimes in antibiotic-mediated antagonistic communities have uncovered mechanisms responsible for *de novo* assembly of diverse microbial communities with higher-order interactions, but such results are qualitatively different from what is possible when mutation is rare (38, 47). What is needed, then, are studies of community assembly in models where a large number of species interact in continuous phenotypic space, are capable of rapid mutation, and engage in both direct and higher-order interactions.

Here, we integrate three major themes concerning the dynamics of microbial communities structured by antagonistic interactions, specifically higher-order interactions, a continuous view of microbial trait space, and an eco-evolutionary perspective on community assembly wherein evolution can happen rapidly. This integrated perspective allows us to explore mixed, rather than strict, patterns of dominance among a large number of biologically realistic species (hundreds to thousands; constrained only by computational power) to study microbial communities of toxin/antibiotic producing, sensitive, and resistant species. Within an agent-based modeling framework, we use a continuous species parametrization model with a wide range of mutational sizes to explore spatio-temporal community assembly, diversity, and stability (see Figure 1 and methods). Parameters are defined at a global level and are divided into two categories: spatial (which define rules and properties of spatial interactions) and species-level (which define the toxin interactions in the species space). The continuous (phenotypic) species space generalizes intra-species relationships of toxin production, vulnerability, and resistance that arise randomly in populations through mutation and interspecies relationships.

**Figure 1:**
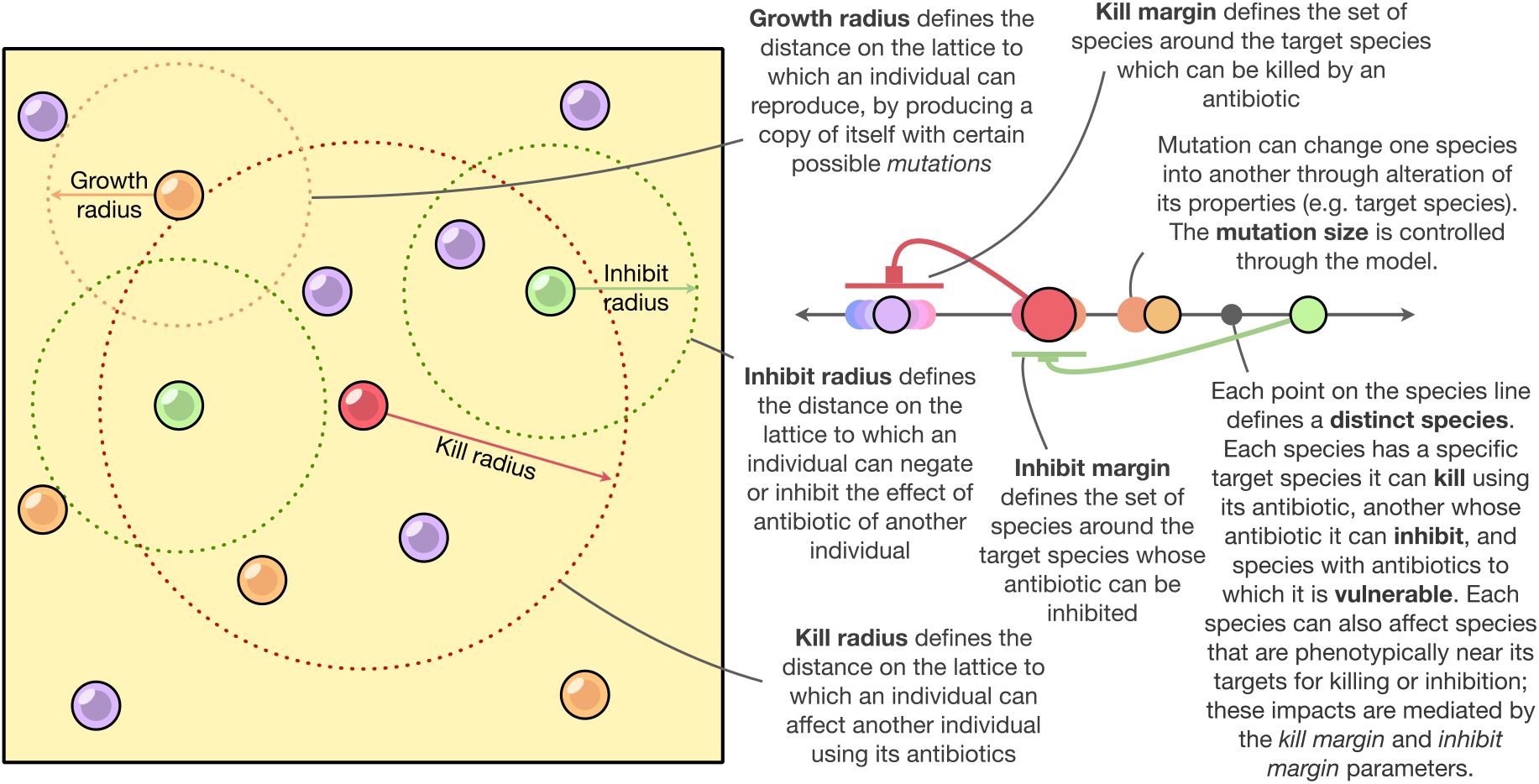
A conceptual representation of the model, with descriptions of all the parameters involved. See Methods for details about implementation.

We find that species-level properties, such as constraints on higher-order interactions among species, affect the spatial structure of microbial dynamics more than spatial factors like mobility and the effective distance over which individuals kill or inhibit each other. In contrast, nonspatial metrics of community diversity reflect complex interactions among species-level and spatial parameters, especially mobility and mutation size, plus community formation time (the time it takes to form a stable community; see 38). Our attention to community formation time, which provides an opportunity for characterizing transient dynamics rather than just focusing on stable states or the long-term maintenance of diversity (15, 39, 47), is intentional. It turns out that community formation time is a very good predictor of diversity dynamics, but this time duration cannot be estimated well from other parameters, pointing at emergent complex properties of community assembly.

## 2. Results

After running 10.49 million simulations across a broad range of parameter combinations, we observed strong differences in spatial heterogeneity among runs. We classified this behavior into three major categories depending only on the observed level of spatial disturbance (low, medium, and high disturbance; Figure 2, Methods). These three categories featured both between- and within-category differences in community spatio-temporal dynamics, but exhibited similar disturbance regimes within categories (Figure 2). The continuous trait axis and the mutable nature of traits together allow for the simulation and visualization of hyper-diverse communities, contrasting with the small, fixed number of discrete, immutable species in previous works (see 6, 20, 27).

**Figure 2:**
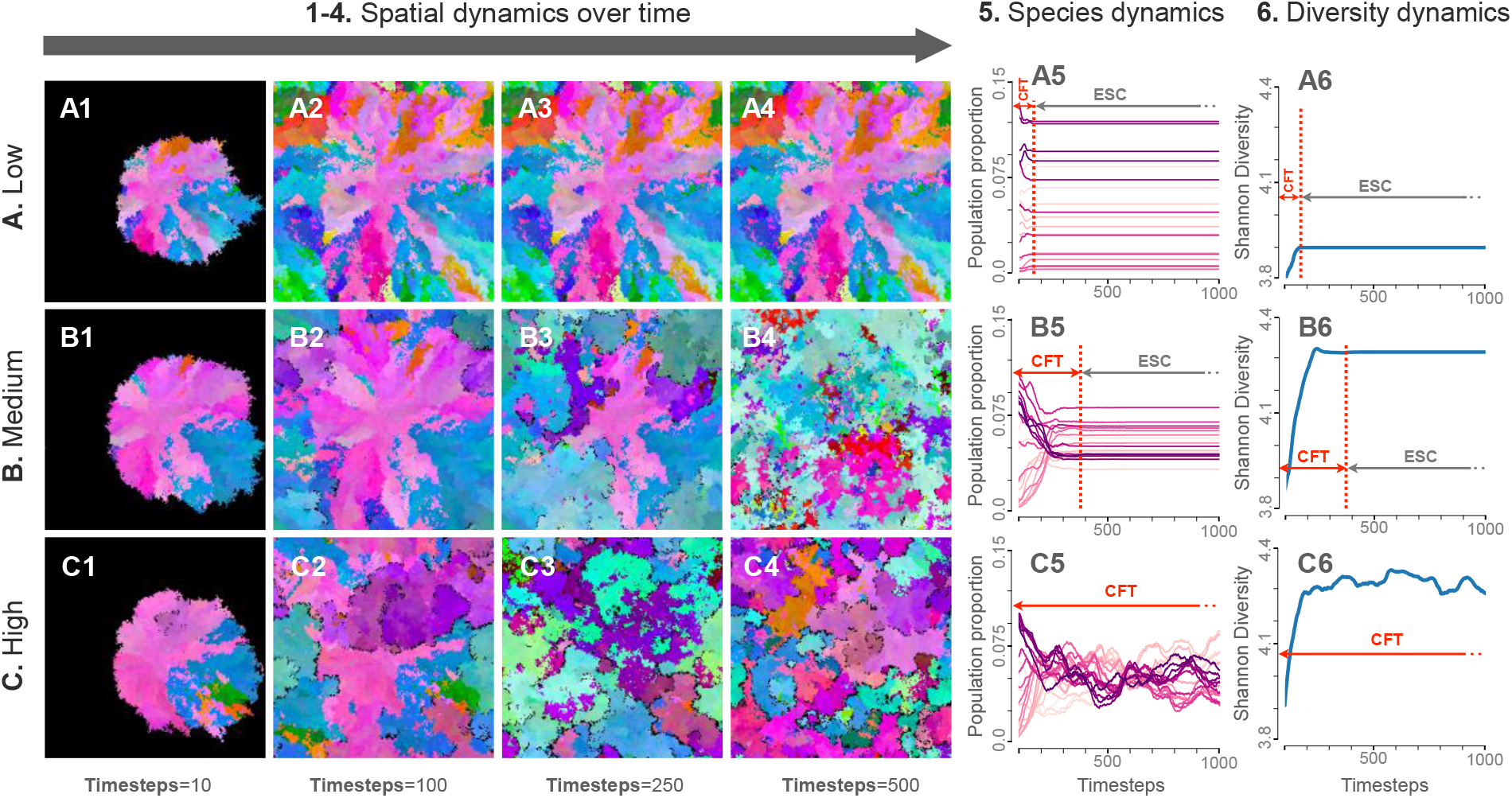
System dynamics in three example conditions of initial conditions pertaining to different regimes of spatial disturbance behavior (Low, Medium and High) (rows A-C). Snapshots of each simulation at different timesteps (10, 100, 250 and 500) are shown in columns 1 through 4. The respective species dynamics (where different colors represent different species ‘bins’, see Methods) and diversity dynamics are represented in columns 5 and 6, respectively. In columns 5 and 6, the time taken for the simulation to stabilize (i.e., the community formation time or CFT) is denoted with a vertical red dotted line. The community existing after the system has reached the CFT is termed an eco-evolutionary stable community (ESC).

The time that each simulation took to stabilize (i.e., to form a stable community) is termed the community formation time (CFT; see Methods and 38), and the community that thus assembled is called an eco-evolutionary stable community (ESC). The incorporation of mutations allows for an exploration of community assembly, transient dynamics (such as CFT and its relation to community properties over time), and complex spatial patterns that was not possible using previous frameworks (Figure 2) (20, 38).

To understand how parameters relate to the spatial disturbance categories, we performed a random forest classification using the six model parameters (*kill radius*, *inhibit radius*, *growth radius*, *kill margin*, *inhibit margin* and *mutation size*). The model had an out-of-bag error of 10.2% and identified *kill margin* and *inhibit margin* as the best predictors of the disturbance classification (Figure 3A). Note that *kill margin* and *inhibit margin* are global species-specific parameters.

**Figure 3:**
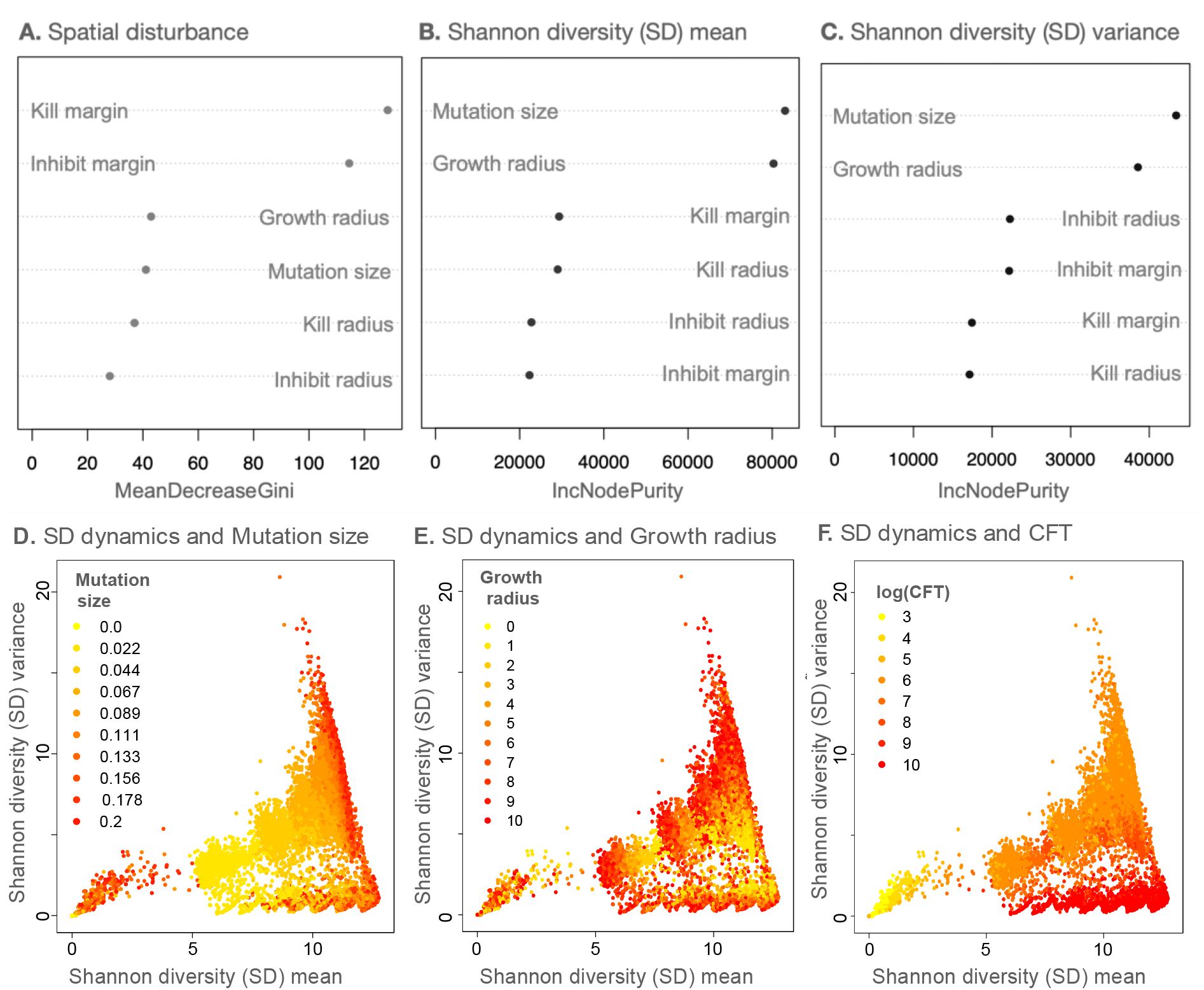
Dependence of model dynamics on different parameters and community formation time (CFT). (A) shows the variable importance score for all the parameters from a random forest classification (with 10,000 trees) on the three categories of spatial disturbance (OOB=10.2%). (B) and (C) denote the variable importance scores from random forest regression models (with 10,000 trees) for Shannon diversity (SD) mean and variance respectively. (D)-(E) show the SD mean-variance plots overlaid with *mutation size*, *growth radius*, and community formation time (CFT) values of the respective simulations. For (A)-(C), IncNodePurity (used in regression random forests) and MeanDecreaseGini (used in classification random forests) refer to how well a parameter explains the prediction variable in question.

The next most important parameter was *growth radius*, which is a proxy for mobility in our model (Figure 1), followed by *mutation size, kill radius*, and *inhibit radius*. These findings showed that, except for *growth radius*, other spatial parameters played a minor role in determining the structure of spatial disturbance in the model systems.

To assess non-spatial metrics of community assembly, we performed a random forest regression on the Shannon diversity (SD) mean (i.e., mean Shannon diversity (SD) for the period from timestep=0 to the CFT) and the associated SD variance using the same six simulation parameters (and an associated species bin size of 0.05 for discretizing species; for more details on species binning, see Methods). That random forest model explained 68.4% of the variance for the SD mean and 70.2% for the SD variance. *Mutation size* and *growth radius* were the two best predictors of both outcome variables (Figure 3B, 3C). Among the less important parameters that affect SD mean and variance, we note that *kill margin* and *kill radius* play a larger role in determining SD mean whereas inhibition processes better predict SD variance (Figure 3B, 3C, S1).

From a naïve interpretation, one would expect *mutation size* to be a good predictor of increasing diversity, as a higher *mutation size* allows for the possibility of greater variation in microbial phenotypic space and therefore the binned SD would be expected to be higher. However, after overlaying the plot of SD mean versus variance with the respective *mutation size* values, the picture is largely muddled (Figure 3D). We saw even more mixed results after overlaying the growth radius values on the SD mean-variance plot (Figure 3E). Overlaying other less important model parameters on the SD mean-variance plot showed no straightforward pattern at all (Figure S2).

Interestingly, upon overlaying the SD mean-variance plot with the values of the respective CFTs, a stronger pattern emerged. Note the clustering by color gradient of the points due to CFT in SD mean-variance space (Figure 3F). Although CFT is not a parameter, it is an emergent outcome of the community assembly process and exploring it further provided insights into the diversity formation process itself.

To formally detect clusters in the SD mean-variance plot with respect to CFT, we used k-means clustering optimized for the number of clusters using multiple methods (Methods) and identified four clusters (Figure 4A, S3). These clusters pertain to (1) Low SD mean and SD variance, (2) High SD mean and SD variance, (3) High SD mean and medium SD variance and, (4) High SD mean and low SD variance, in order of increasing values of CFT.

**Figure 4:**
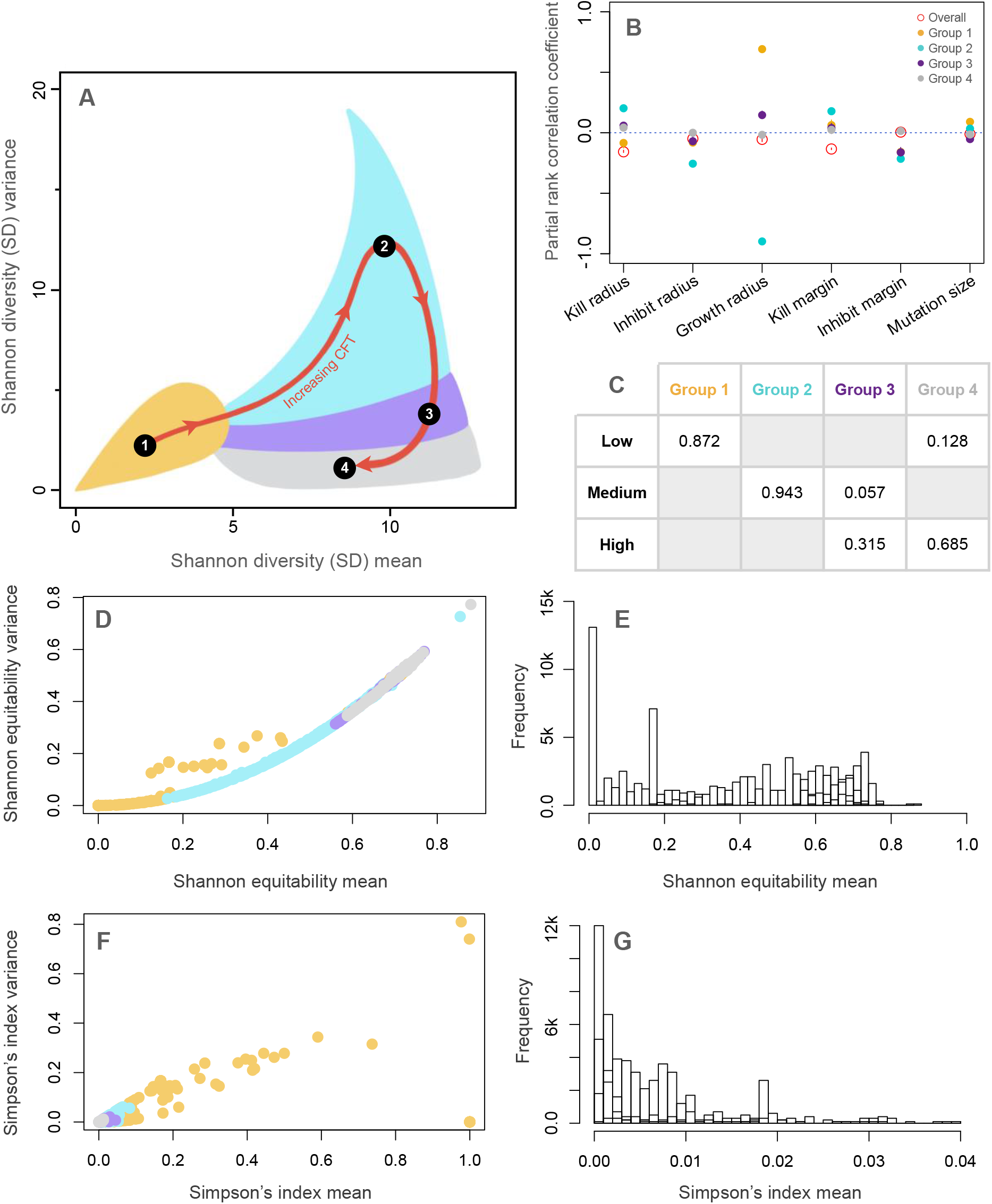
Effects of Community formation time (CFT) on the spatial diversity dynamics. **(A)** depicts the four optimal clusters (groups) of SD mean-variance space found using k-means clustering (see Figure S3). **(B)** shows the results from the partial rank correlation coefficient (PRCC) analyses of the four groups and the overall data. **(C)** shows the proportion of spatial disturbance regimes present in different groups. **(D)** and **(F)** show the distribution of the values of mean and variance of Shannon Equitability and Simpson’s index, respectively, for a smaller subsample of the runs, with points colored by groups from **(A)**. **(E)** depicts the group-wise histogram of Shannon equitability and **(G)** the histogram of Simpson’s index (for clarity, only the range between 0 and 0.04 is shown).

On performing random forest regression predicting CFT using the six-parameters for the whole dataset, we found a low explanation of variance (22.3%), with the *kill parameters* (*kill radius*, and *kill margin*) being the strongest predictors. However, upon performing the analysis separately on the four CFT groups, a stronger pattern emerged (Figure S4). For group 1, the regression explained 30.2% of the variance with *growth radius* being the strongest predictor, and other variables playing a smaller part, but nevertheless the second and third best predictors are *kill parameters* (Figure S3A). Group 2 regression explained 90.6% of the variance with again *growth radius* as the strongest predictor with other parameters making almost no contributions (Figure S4B). For group 3 and 4, the regression explains 27.2% and 6.0% of the variance respectively, with *growth radius* again having most impact on prediction (Figure S3C,D). This pattern of results is unusual because the overall regression did not show any strong dependence on *growth radius* (Figure S3E).

To explore this dilemma further, we performed a partial rank correlation coefficient analysis on this data pertaining to CFT (Figure 4B). We found that the sign and strength of the dependence of *growth radius* for CFT groups vary widely; hence, the random forest regression on the whole dataset misses this signal. *Growth radius* had a strong positive PRCC with CFT in group 1, a very strong negative PRCC with CFT in group 2, a weakly positive PRCC in group 3 and a negligible one in group 4. Other parameters had relatively smaller PRCC values as compared to *growth radius* values for group 1 and 2. *Kill radius* and *kill margin* had overall higher PRCC values than the other parameters and thus were identified by the random forest algorithm (Figure 4B, S4E). Moreover, on calculating PRCC of parameters on SD mean and SD variance, we found that the general values are similar for both SD metrics, but what differentiates SD mean and SD variance values the most is *growth radius* (Figure S1).

To link the dependence of diversity dynamics on CFT with the spatial disturbance regimes, we compared what proportion of the three spatial disturbance regimes (low, medium, and high) fall into the four CFT-classified groups from SD mean-variance space (Figure 4C). We found that the low disturbance regime was most associated with group 1 and, to a much lesser extent, group 4. The medium disturbance regime was primarily dominated by group 2, with a small contribution from group 3. Lastly, the high disturbance regime was dominated by groups 3 and 4. These results can be interpreted in the light of Shannon equitability and Simpson’s index values for the simulations (Figure 4D-G). Transitioning from Group 1 to 4 reveals an increase in mean Shannon’s equitability (Figure 4D,E) and a decrease in mean Simpson’s index (Figure 4F,G). One interpretation here is that higher equitability among species (i.e., higher values of Shannon equitability), which can occur when comparatively few species dominate the community (i.e., lower Simpson’s index), results in simulations that take longer to equilibrate (i.e., longer CFTs). For example, communities in Group 4 take longer to equilibrate (longer CFTs) as the species in those communities are more nearly equitable, with comparatively fewer species whose interactions exercise control over the community. In contrast, Group 1 has low equitability, implying a higher level of community dominance by some players, and therefore, those simulations settle down quickly (low CFTs). Collectively, these results highlight some important, and novel, connections between temporal spatial patterns and diversity dynamics, especially through the lens of transient phenomena (see 52, 53).

All the results in this section were performed using a species bin size (for discretization of species for diversity metrics; see Methods) of 0.05, but we repeat the analyses at three other reasonable bin sizes (0.03, 0.07 and 0.09; see Methods) and find almost no changes in the structure of the results and their interpretations (see Figure S5) in any of the cases. The space of SD mean and variance shrinks in scale (Figure S5C) as the bin sizes increase, which is expected because increasing bin sizes results in fewer distinct species for analysis. However, the categorization of spatial disturbance remains unaltered because binning does not affect the actual dynamics of the model, but only how we quantify those dynamics through discretization. Therefore, the dependence of CFT on parameters remained unaltered (Figure S5D). The categorization of SD mean and variance into groups, their relation to spatial disturbance regimes, and their dependence on parameters also varied little (Figure S5A,B,E).

## 3. Discussion

A continuous species model with simple rules for interactions based on antibiotic production, inhibition, and vulnerability can produce a wide array of complex dynamics and generate hyper-diverse, persistent communities. Despite enormous heterogeneity in behavior, certain general patterns in the assembly and stability of these hypothetical microbial communities stand out.

The diversity patterns seen across time were expressed in the form of mean Shannon diversity (SD) and its associated variance. The phase space of these two metrics shows a non-linear relationship between the metrics. Random forest regressions revealed that *mutation size* and *growth radius* were the best predictors of mean SD diversity and its variance (Figure 3B,C). In other words, the mutation size and mobility of species jointly determine how diversity dynamics play out in these theoretical communities.

Mobility controls diversity dynamics in numerous previous studies (6, 20, 27, 54). In particular, small amounts of mobility enhance diversity, whereas large amounts of mobility jeopardize it (6, 54). Reduced mobility results in spatially structured populations where co-existence is easier to maintain, as seen in experimental works on competing *E*. coli strains (26, 55, 56). Recent work has shown that even in cases of high mobility (i.e., well-mixed communities), one can observe co-existence if higher-order interactions, such as antibiotic production and degradation, are considered (20). This is the case for *in vivo* experiments with bacterial colonies in the intestines of co-caged mice; these systems can be considered locally well-mixed and have high levels of coexistence (30).

High mutation rates, and processes such as horizontal gene transfer (HGT), have been long known to affect diversity in experimental microbial populations, especially during the initial phase of community assembly (57, 58). But cyclic dominance models have tended to overlook the role of mutations and HGT in coexistence. Such studies have focused instead on detailed understanding of a small number of discrete species under a low mutation regime (47). Recently, studies have focused on understanding the effects of high mutational regimes in community assembly, which better represent microbial systems (38), and can generate frequent non-cyclic interactions (40). Our efforts have considered an array of mutational regimes and characterized the importance of non-cyclic patterns of interactions in the assembly and maintenance of microbial communities.

*Mutation size* and *growth radius* (a proxy for mobility) were good predictors of SD mean and SD variance but yielded no discernable patterns in the phase space (Figure 3D,E). In contrast, community formation time (CFT) partitioned the space cleanly into four regions with distinct properties (Figure 3F, 4A). Communities that have low diversity (SD mean) and small fluctuations in diversity (i.e., low SD variance) (group 1) assemble quickly, and mobility enhances diversity and co-existence in this scenario (Figure 4A,B). In contrast, mobility negatively affects diversity and coexistence for communities with high diversity (SD mean) and larger fluctuations (high SD variance) (group 2), and these communities assemble slower than group 1. For communities that take a long time to stabilize (groups 3 and 4), diversity is generally high (SD mean), but it fluctuates less and mobility has a negligible effect.

Mutation sizes were weakly correlated with CFT, as observed previously (38) but exhibited substantial variation in trends due to interactions with other model parameters. For low SD mean and variance (group 1) and high SD mean and variance (group 2), *mutation size* positively affects CFT (Figure 4B). This implies that communities with larger mutations take longer to stabilize, but only when the diversity outcome corresponds to these two groups (Figure 4B). If SD mean is high but SD variance is low, mutation affects CFT negatively, i.e., higher mutation helps communities converge faster (Figure 4B).

Community spatial structure is also important for coexistence (54). However, rather than focusing on the final spatial structure, we examined the degree of spatial disturbance that occurred throughout the assembly process, which provides insight into the extent of mixing/migration that might occur in these communities. The categorization of spatial disturbance regimes was best predicted by species-level parameters (*kill margin* and *inhibit margin*), rather than by other spatial parameters in our model. This is somewhat surprising, but higher-order interactions are known to influence spatial patterns of coexistence (20). Importantly, we found a strong correspondence between the regimes of spatial disturbance and the groups produced by CFTs. The communities that formed quickly (group 1) tended to have low spatial disturbance. (Figure 4C). Communities that took more time to stabilize and ended up with high diversity through large overall diversity fluctuations (high SD mean and SD variance, group 2) have primarily a medium level of spatial disturbance. The communities that took longest to stabilize were characterized by high diversity (SD mean) and lower average fluctuations in diversity (SD variance) (groups 3 and 4) but have larger spatial disturbances.

All these observations point towards a complex interplay of mutation and mobility, which affects the assembly time of a community (CFT), and in turn controls the diversity of the assembled community. Although mobility (or more broadly, dispersal) is known to enhance diversity in ecological communities in both theoretical (59, 60) and experimental (61, 62) settings, especially at local scales (63), we found that mobility is linked to diversity only in communities with short and intermediate CFT (i.e., those that equilibrated relatively quickly). Mobility enhances diversity and coexistence in Group 1, where a subset of dominant (high relative abundance) species appear to set community dynamics. In this capacity, dispersal would appear to act in the classic disruptive fashion, permitting coexistence where it would otherwise not occur (64, 65). In contrast in Group 2, where communities are characterized by both high diversity and high fluctuations, mobility has a negative effect on diversity and coexistence, in keeping with the capacity for dispersal to homogenize otherwise diverse systems (66, 67). This dichotomy, together with the absence of an important role for mobility in Group 4, where communities are characterized by long-term nonequilibrial dynamics, offers an intriguing target for future integrative research.

The mechanistic light shed upon these patterns by studying the Shannon equitability index (which measures the distribution of relative abundance of species in a community) and Simpson’s index (which measures the distribution of ‘dominance’ of species in a community) is also worth mentioning. The more equitable a community (higher Shannon equitability), the lower the chance of a particular species dominating the interactions (lower value of Simpson’s index) and therefore, the longer it takes for the community dynamics to stabilize (longer CFTs), as seen in the behavior of the four groups (see Figure 4D-G). The importance of species dominance to the maintenance of diversity in this model, which is structured via non-resource based antagonistic interactions, is intriguingly similar to the critical role that species dominance plays in both real and theoretical communities structured via resource-based competition (68–71).

Collectively, these results demonstrate the rich spatiotemporal dynamics that are possible when large numbers of microbial species with limited but heterogeneous rules for aggressive, inhibitory, and vulnerable interactions live in a common space. Although these findings are focused primarily on non-resource-based antagonistic interactions between microbes, such dynamics may also be relevant for coral communities and other spatially structured systems featuring diverse types of interspecific interactions (see 72–74).

Our results also point towards the important yet often-overlooked role that transient dynamics play in the behavior and structure of ecological systems (see 53). Usually, models of ecological dynamics use asymptotically stable behavior or values at stability to explore patterns in the system (53). Admittedly, we have followed this approach in our exploration of how different parameters affect microbial communities’ long-run behavior (i.e., after CFT is reached). In addition, however, we also focused on categorizing spatio-temporal disturbance regimes and connecting these transient phenomena to system diversity. Great opportunities exist for future work investigating temporal diversity dynamics and how they relate to spatial heterogeneity and system processes. Such investigations can shed light on how transient microbial dynamics and assembly history affects the species diversity seen in microbial communities (75).

Even though our model introduced the use of continuous axes for species traits and interactions, we note that the *post-hoc* binning procedure creates a bridge back to the matrix/network framework that has characterized community ecology studies for decades (2, 4, 13, 14, 21, 23). In this case, however, a multi-layered network perspective would be necessary to accommodate the different kinds antibiotic-mediated interactions (production, inhibition, and vulnerability). Future work could also incorporate other important forms of microbial interactions (e.g., mutualism, cross-feeding, resource competition) into the continuous species framework we have developed. Such investigations would allow exploration of how the interplay between resource-based and antagonistic interactions jointly shape diversity dynamics. However, to do this we need to understand the interrelationships between resource-based and antagonistic interactions across species (see 11, 76, 77). Such investigations could be highly beneficial by providing a more complete view of how higher-order, intransitive interactions shape community assembly, stability, and diversity in natural systems.

## 4. Model

### 4.1 Model Description

Here we outline an algorithmic presentation of the model. The parameters of the model and their corresponding symbols are detailed in Table 1. The simulation takes place on a 200 x 200 square lattice with toroidal boundary conditions. This size of this simulation was a tradeoff between being small enough for computational feasibility and large enough to allow interesting dynamics. We found 200 x 200 to be a reasonable range for our computation budget. The model of microbial dynamics on this simulation is split into two parts: the microbial (phenotypic) space and the environment space (see Figure 1). This division of the model is purely for convenience and can easily be reinterpreted where the microbes and the environment are not differentiated. For the microbial phenotypic space, each cell in the lattice represents a single ‘variant’ of microbe and is parameterized by a stateful three-dimensional species vector ***R*^3^*ϵ*** {Ø, [**0, 1**]**^3^**}, where Ø represents a cell that is unoccupied and [0,1] is a bounded real value that defines the parameters of any occupants. The three elements of the species vector are the species’ vulnerability (*S_ν_*), the species’ inhibition value (*S_i_*), and the species’ antibiotic production value (*S_a_*). In the microbial space, neighboring cells do not interact with each other directly, but instead affect the environment space which then affects the microbial space. The environment space is stateless and instead always resets to the most recent effects from the microbial space, or in other words, no antibiotics remain in the environment between timesteps. This can just be interpreted as diffusion on a less granular timescale. The environment space is parameterized by a two-dimensional environment vector ***R*^1^*ϵ*** {Ø, [**0, 1**]}, where its single element represents the antibiotic occupying that cell space (*C_a_*), but again, the cell can also contain no antibiotics, represented by Ø. Please note that we choose a phenotypic microbial space that maps genotypic changes into phenotypic traits. Doing this in a meaningful way is a very difficult problem, and this is especially true for bacterial toxin-based antagonisms, where the basic nature of the interactions remains at a preliminary level of research and discovery (11). In our case, this means that the microbial space is not uniformly mapped to the size of the genetic changes. In this simple model, however, this does not create a problem as we only deal with ‘expressed’ phenotypic changes.

**Table 1:** Brief description of model parameters

| Parameter | Type | Definition |
| --- | --- | --- |
| *Species antibiotic production value* ($S_a$) | Species-level,<br>Species-specific | Defines what species are vulnerable to the antibiotic that this individual produces |
| *Species vulnerability value* ($S_v$) | Species-level,<br>Species-specific | Defines what antibiotics (and hence species) this individual is vulnerable to |
| *Species inhibition value* ($S_i$) | Species-level,<br>Species-specific | Defines the species whose antibiotics this individual can inhibit |
| *Kill radius* ($K_r$) | Spatial, Global | Maximum spatial distance of diffusion of the antibiotics produced by an individual |
| *Inhibit radius* ($I_r$) | Spatial, Global | Maximum spatial distance of diffusion of the inhibitors to antibiotics of produced by an individual |
| *Growth radius* ($G_r$) | Spatial, Global | Maximum spatial distance at which an individual can reproduce by producing a copy of itself with certain possible mutations |
| *Kill margin* ($\epsilon_a$) | Species-level,<br>Global | The total span centered around the specified *Species antibiotic production value* ($S_a$), which can be affected/killed by the antibiotic of a given species |
| *Inhibit margin* ($\epsilon_i$) | Species-level,<br>Global | The total span centered around the specified *Species inhibition value* ($S_i$), whose antibiotic can be inhibited by a given species |
| *Mutation size* ($\mu$) | Species-level,<br>Global | Maximum value by which the species-specific parameters can be additively altered during reproduction/copying |

At fixed-interval time steps, the state of the microbial space is updated synchronously in four stages. First, an antibiotic release step occurs, which simulates the process of microbes diffusing antibiotics into the environment. Here each cell’s antibiotic value (*C_a_*), in the environment space is randomly assigned a species’ antibiotic value (*S_a_*) of one other cell from the microbial space that is within the antibiotic radius (*K_r_*). Note that rewrites of the environment cell are possible during this step, so the order of cell updates is randomized. Second, an inhibition step occurs which handles any inhibiting effects of the antibiotics in the environment space. In this step, for every cell in the microbe space, an inhibition radius (*I_r_*) around it is checked for other cells in the environment space which contain an antibiotic value (*C_a_*) such that the inhibition value is within a marginal range of the antibiotic value, *I_ν_*-*ϵ_i_* ≤ *C_i_* ≤ *I_ν_* + *ϵ_i_*. If this is the case, the cell in the environment space antibiotic value (*C_a_*) is made empty, Ø. Third, a kill step occurs where we iterate over every cell, comparing the antibiotic value (*C_a_*) in the environment space and the individual vulnerability value (*S_ν_*) in the microbial space. If the individual vulnerability value (*S_ν_*) is within the range of the antibiotic value (*C_a_*). *C_a_* – *ϵ_a_* ≤ *S_ν_* ≤ *C_a_* + *ϵ_a_* then the individual is killed and the cell in the microbial space is made empty, Ø. Fourth and last, a growth step occurs, where for every cell in the microbial space that it is not empty, another cell in the microbial space that is within that cell’s growth radius (*G_r_*) is randomly selected. If the selected cell is empty, the species vector of the growing microbe cell is copied to the selected cell. The copied parameters are also further modified according to a mutation size (*μ*). This modification is a non-correlated additive perturbation of the species’ vector by a uniform value sampled from *U*(–*μ*,*μ*) and the value is restricted in the range [0,1] by adding or subtracting 1 if it is outside this range. This ensures that all species have equal likelihood of interactions with other species by putting the species parameters on a three-dimensional toroid. From a theoretical perspective, this growth can be seen as a model of growth probability, where the probability that an individual will reproduce is equal to *E*(*C*)/*F*(*C*) where *E*(*C*) is the number of empty cells within radius *G_r_* of the current *C* and 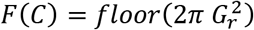 is equal to the total number of cells within radius *G_r_* of cell *C*.

### 4.2 Model Assumptions

There are three major assumptions/rules underlying our model implementation. The first one is that we keep all spatial (environment space) parameters, mutation size and margin of interactions constant in a given single run while mutations affect and alter species identity parameters only (*S_a_*,*S_ν_*,*S_i_*). We assume this because properties like kill radius and inhibition radius are typically affected by diffusion properties of the toxin compounds, which are of similar/comparable physical properties in a given class of molecules (but can vary across classes of toxins; see 11). Hence, for computational purposes, we keep these parameters constant over a run, along with the generality of their interactions (controlled through the margin parameters (*ϵ_a_*,*ϵ_ν_*,*ϵ_i_*)). We keep the mutation size constant for simplicity and for comparisons across different mutational regimes.

The second assumption pertains to the structure of species traits in the microbial space. The phenotypic space, as mentioned, will be non-uniformly mapped to genetic changes in species (i.e., no correlation exists between distance on the phenotypic space along any axis and the genetic change from which it results). Given the dearth of empirical information on the structure and dependencies of these traits (e.g., whether they are independent, or constrained; see 11), we assumed an independent, normalized space of microbial traits for simplicity. This assumption can be restructured once additional research clarifies the nature of the dependencies between genotypic and phenotypic changes.

Lastly, we build our model to attend only to toxin-based antagonistic interactions. Little evidence is available to characterize the relationship between antagonistic interactions and other forms of microbial interactions (e.g., mutualisms, resource competition) across species. Therefore, our aim here was to build a ‘base’ model that allowed for the investigation of antagonistic interactions with more nuance and across more dimensions of complexity than was possible in previous work on the subject. As more information becomes available about the interplay between various types of microbial interactions, these can be incorporated together in future work.

### 4.2 Model Implementation

We implemented the simulation in *C*++ using the *SFML* graphics library for rendering. It is possible to parallelize the updating of sub-steps of the cells due to the synchronous nature of the model, but we chose to use a simpler sequential algorithm that looped through each cell one at a time. We did, however, parallelize the running of multiple simulations when sweeping parameter spaces. This allowed us to reduce the total computation time required by two orders of magnitude. Each simulation was run for 2000 timesteps or until stability/stagnation (also termed the Community Formation Time (CFT)). Stagnation/stability was determined by comparing the state of all cells between two consecutive time steps, and if it remained unchanged for a total of five-time steps, the simulation was terminated. This helped to quickly remove a large number of simulations that terminated early due to poor parameter settings. Because the simulation was so large, collecting per-cell data across many parameters was not practical. Alternatively, we chose to compute population-level statistics online during each simulation using Shannon Diversity mean and variance (over time) and then from the analysis of this lower-granularity data, we selected individual parameter settings to explore population-level statistics. Further, we also built a qualitative application to allow quick classification of simulations, assigning them to a class based on their spatial heterogeneity/disturbance dynamics. Simulations were assigned to one of three classes according to how the spatial dynamics evolved (Fig. 2).

To track population-level statistics, it was necessary to discretize the continuous species space. *l*-discretization (or *l*-binning) assigned individuals to the same species if the three parameters of those individuals had the same integer values after performing a binning transformation on them like so: 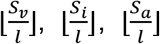. When recording the population-level data, we stored the number of individuals per species every time step, using a standard 0.05-discretization. Although the choice of 0.05-discretization is arbitrary, we emphasize that the binning only affects the resolution of the statistical analysis and has no influence on the dynamics of the model itself. We found 0.05 to be a small enough value for which the simulation statistics did not blend away, but not too small such that there were no measurable statistics (see Data Analysis, Section 4.3). For comparison, we repeated the complete set of analyses on system properties at three other reasonable species bin sizes (0.03, 0.07, 0.09) (see Fig. S5 for more details).

For every parameter setting, we repeated the simulation ten times, seeding the pseudo-random number generator with a unique value for each repetition. Every simulation began by assigning every cell vector the all-empty state (−1, −1, −1, −1, −1) except the center cell which is assigned a random value in the range [−1,1] for the first three elements of its vector. This initial cell represented a randomly created individual. One potential issue was that if the center cell was assigned a random value such that it was vulnerable to its own antibiotic, it would kill itself and the simulation would terminate immediately. Although this was possible, it was rarely the case, and it did not significantly affect our results due to our use of multiple runs per parameter setting.

### 4.3 Data analysis

We ran a total of 10.49 million runs of our model with various parameter initializations, and performed data analysis with *Python 3.7* and the *R* statistical language. For these runs, we obtained a manual-classification of the size of the spatial disturbance (low, medium and high). The simulations which stabilized quickly (fewer than 20 time-steps) with no further changes in spatial patterns and the ones that had (more-or less) stabilized by 20 time-steps, except for a small number of point fluctuations in the grid, were categorized as the ‘low disturbance’ regime (Figure 2 A1). The simulations which, had not stabilized after 20 time-steps but exhibited small waves of diversity replacements which diminished over time were termed as ‘medium’ disturbance. All medium disturbance simulations stabilized before 500 time-steps (Figure 2 B1). The remaining simulations did not stabilize even after 500 time-steps, nor had reduced amplitude of waves of diversity replacements; these were categorized as the ‘high’ spatial disturbance regime (Figure 2 C1).

We also calculated the mean Shannon diversity, Shannon equitability, and Simpson’s index over time (until a community stabilized, i.e., reached the community formation time, CFT), and the variance in each of the three metrics over the same time period. The disturbance classification, SD mean, and SD variance were our three outcomes of interest and are the focus of the results. We later connect the results to the values of Shannon equitability and Simpson’s index.

To understand how parameters affect diversity dynamics, we determined the partial rank correlation coefficient (PRCC) of various parameters with respect to the three outcomes of interest, using the *sensitivity* package (78) in R. We also performed Random Forest (RF) regressions and classification, using the *randomForest* package (79) in R, to identify which parameters were the strongest predictors of the patterns in predicting the outcomes of interest. We optimized the number of parameters available for splitting at each tree node in the RF using out-of-bag error (OOB) (79). Please note that the overlay figures (Figure 3 D-F, Figure S2) have been plotted with pruned samples for better visualization.

For clustering values on the SD mean versus SD variance space with respect to CFT, we used k-means clustering in *R* and optimized for the number of clusters using the elbow method, the D index, and the Hubert indices (which all yielded similar results) using the package *NbClust* (80). For each obtained cluster in the Shannon diversity space, we performed a random forest analysis to calculate differences in variable importance scores across these clusters. For each cluster, we also performed PRCC and random forest analyses, in order to delineate the effects that individual parameters had on the different subsets of the diversity space.

## Author Contributions

AS, LF and WFF conceptualized the manuscript; AS and LF performed the simulations, data analysis and visualizations; AS, LF and WFF wrote and edited the manuscript

## 5. Acknowledgements

NSF DMS-1853465 (to WFF) and the University of Maryland supported this work.

## 6. Code and Data Availability

All code and required data to replicate our results are available in the following repository: https://github.com/levifussell/MicroEvo.

