## Supplementary figures for "Higher-order effects, continuous species interactions, and trait evolution shape microbial spatial dynamics"

**This PDF file includes:**

Figures S1 to S5

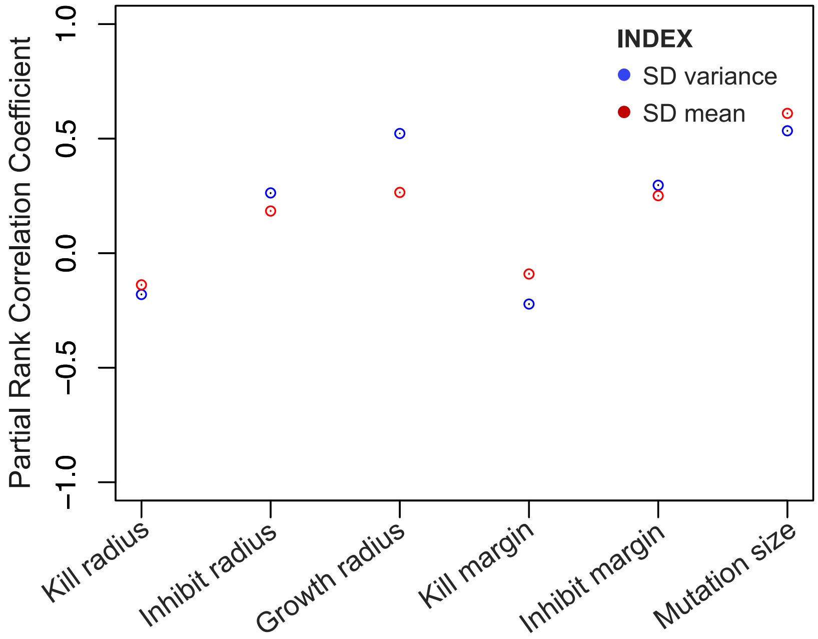


**Figure S1:** Partial rank correlation coefficients (PRCC) of model parameters in relation to the Shannon Diversity (SD) mean and variance. Please note that different parameters have similar values of PRCC for both the diversity metrics, and the greatest difference in the trends of the two metrics was for *growth radius*.

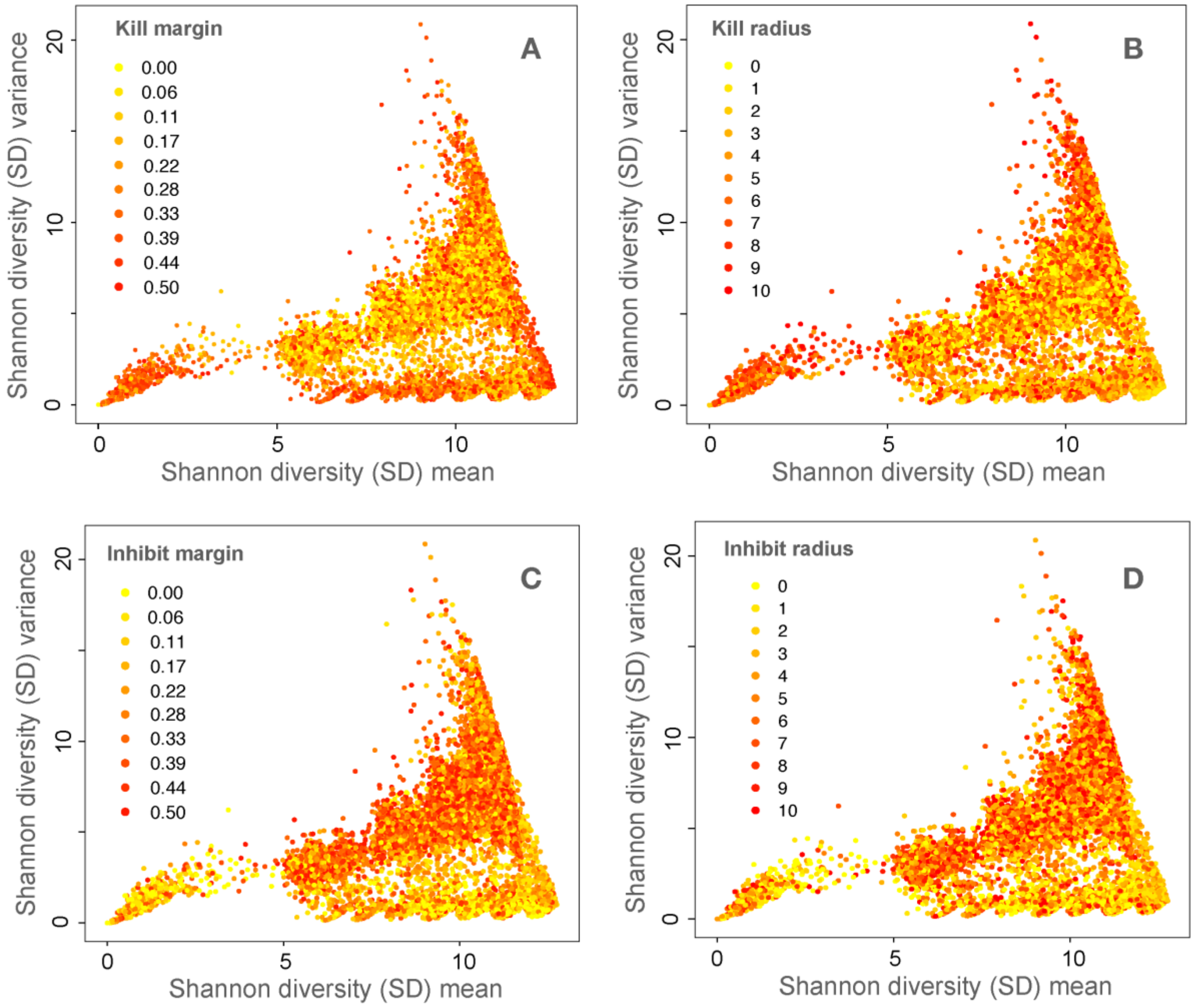


**Figure S2:** Overlaying values of parameters, (A) *Kill margin*, (B) *Kill radius*, (C) *Inhibit margin*, and (D) *Inhibit radius*, over the respective values on Shannon Diversity (SD) mean and SD variance space. No obvious patterns are discernable in how these parameters affect the diversity space as compared to the ones in Figure 3 D-F, where distinct patterns in clustering of the diversity space can be observed.

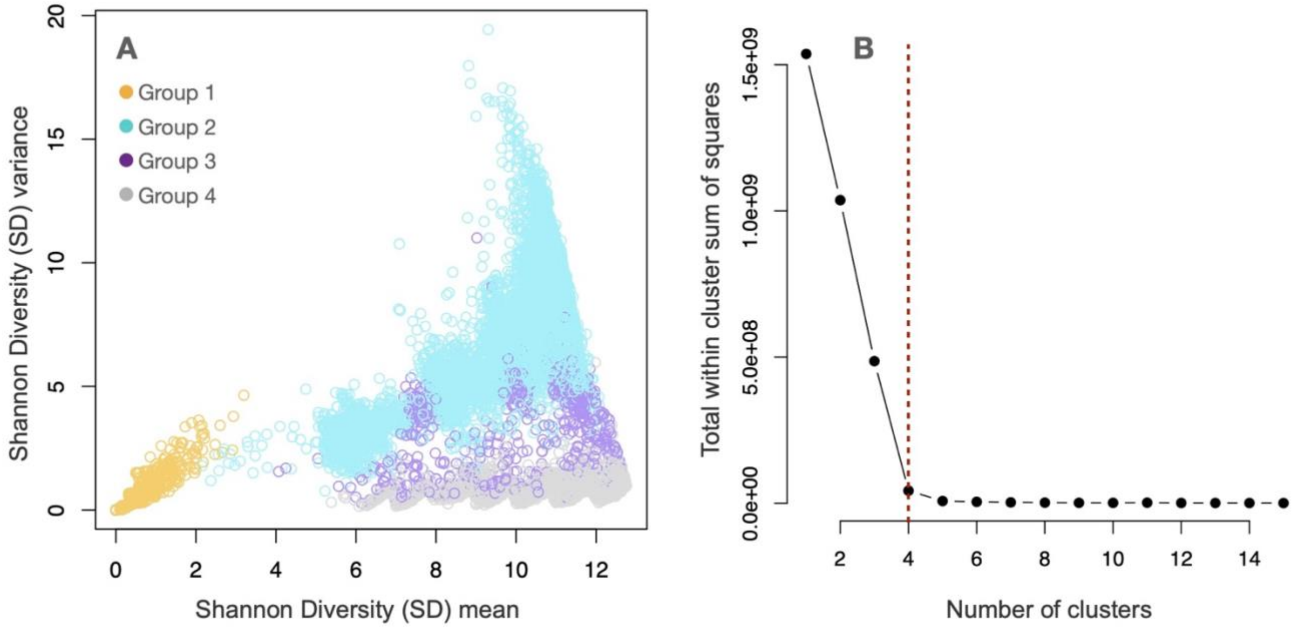


**Figure S3:** (A) k-means clustering of the Shannon Diversity (SD) mean-variance space by the community formation time (CFT) value. (B) shows the optimal number of clusters using the elbow method. We obtained similar results when optimizing clusters using the D index and the Hubert indices.

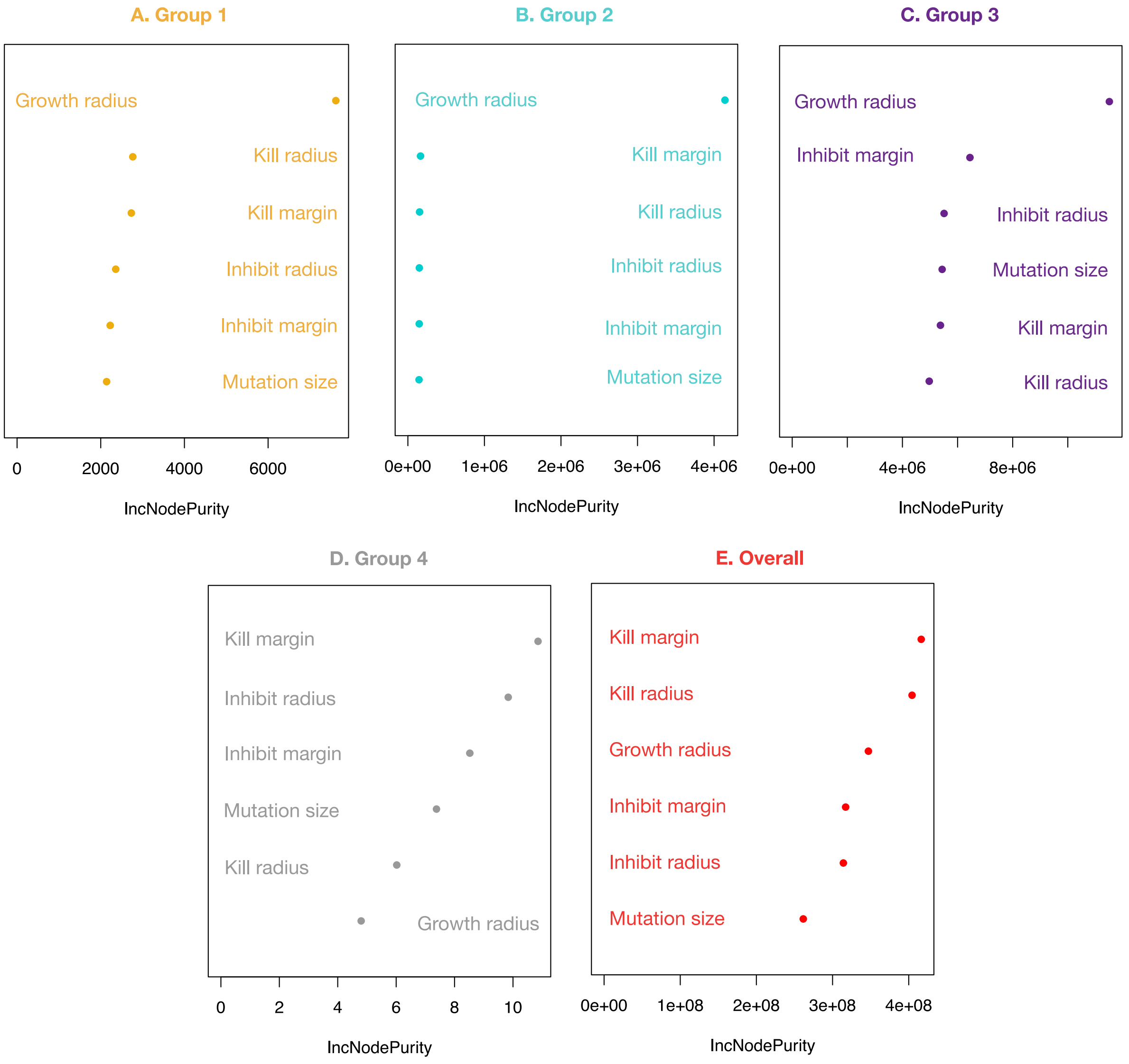


**Figure S4:** Random forest regressions (10,000 trees each with optimization for splits using out-of-bag (OOB) error) for community formation time (CFT) for the four diversity groups clustered using k-means. Variance explained for- Group 1: 30.2%, Group 2: 90.6%, Group 3: 27.3%, Group 4: 6.0%, Overall: 22.3%. Note: IncNodePurity refers to the total decrease in node impurities from splitting on a given parameter, averaged over all trees.

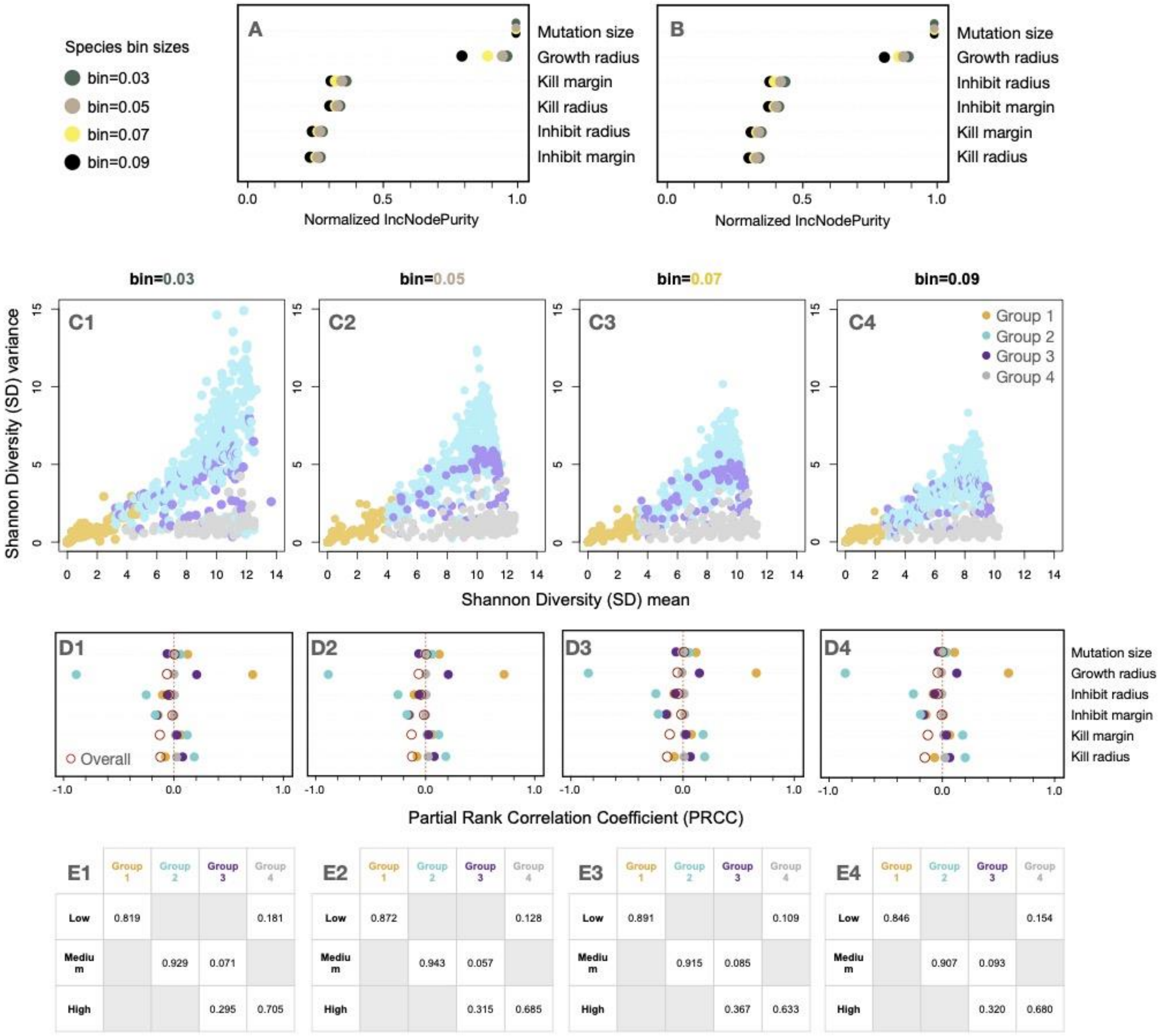


| E1 | Group 1 | Group 2 | Group 3 | Group 4 |
|---|---|---|---|---|
| Low | 0.819 | | | 0.181 |
| Medium | | 0.929 | 0.071 | |
| High | | | 0.295 | 0.705 |

| E2 | Group 1 | Group 2 | Group 3 | Group 4 |
|---|---|---|---|---|
| Low | 0.872 | | | 0.128 |
| Medium | | 0.943 | 0.057 | |
| High | | | 0.315 | 0.685 |

| E3 | Group 1 | Group 2 | Group 3 | Group 4 |
|---|---|---|---|---|
| Low | 0.891 | | | 0.109 |
| Medium | | 0.915 | 0.085 | |
| High | | | 0.367 | 0.633 |

| E4 | Group 1 | Group 2 | Group 3 | Group 4 |
|---|---|---|---|---|
| Low | 0.846 | | | 0.154 |
| Medium | | 0.907 | 0.093 | |
| High | | | 0.320 | 0.680 |

**Figure S5:** Dependence of the results on species bin sizes. (A) and (B) depict the normalized variable importance scores for a random forest regression of Shannon diversity (SD) mean and variance, respectively, at the four bin sizes. Percent variance explained remained between 68-75% for all bin sizes, and all other results were similarly consistent. (The only thing to note is that the dependence on growth radius (a mobility-related metric) weakens with increasing bin size, which may occur because increasing bin size results in greater discretization for a given length scale). (C1-4) depict the k-means clustering of the space of SD mean and variance for the four bin sizes. Increasing bin size collapses the range of SD mean and SD variance of the resultant communities, but otherwise no changes are obvious. (D1-4) depict the Partial Rank Correlation Coefficients (PRCC) of variables on Community Formation Time (CFT), which are consistent across bin sizes. (E1-4) depict how the spatial categorizations correspond to the diversity groups at four bin sizes, and again the results are consistent.
